# The cost of heat waves and droughts for global crop production

**DOI:** 10.1101/188151

**Authors:** Zia Mehrabi, Navin Ramankutty

## Abstract

Heat waves and droughts are a key risk to global crop production and quantifying the extent of this risk is essential for insurance assessment and disaster risk reduction. Here we estimate the cumulative production losses of six major commodity groups under both extreme heat and drought events, across 131 countries, over the time period of 1961-2014. Our results show substantial variation in national disaster risks that have hitherto gone unrecognised in regional and global average estimates. The most severe losses are represented by cereal losses in Angola (4.1%), Botswana (5.7%), USA (4.4%) and Australia (4.4%), oilcrop losses in Paraguay (5.5%), pulse losses in Angola (4.7%) and Nigeria (4.8%), and root and tuber losses in Thailand (3.2%). In monetary terms we estimate the global production loss over this period to be $237 billion US Dollars (2004-2006 baseline). The nations that incurred the largest financial hits were the USA ($116 billion), the former Soviet Union ($37 billion), India ($28 billion), China ($10.7 billion) and Australia ($8.5 billion USD). Our analysis closes an important gap in our understanding of the impacts of extreme weather events on global crop production and provides the basis for country relevant disaster risk reduction.

## Main text

There is now credible evidence that human driven climate change is leading to an increase in the severity and frequency of extreme weather events^1^. However, the agricultural risk associated with these extreme events is not only a function of their frequency: it also depends on whether they occur in key production locations, and how vulnerable production systems are to their onset^2,3^. Despite calls from within and outside the scientific community to determine agricultural risk under extreme weather disasters^2,4,5^, and the identification of heat waves and droughts as key components of this risk^3^, we still know surprisingly little about the impact of disaster events on crop production at the global level. There are at least three agenda setting knowledge gaps that need to be filled. First, previous work on the impact of these events has averaged impacts at a regional level^3^. We need to estimate risk on the country level to align scientific efforts with international disaster risk response profiling initiatives (e.g. INFORM^6^). Second, average per-impact loss estimates^3^, are important for determining stocking requirements for isolated events, but do not give the full picture of risk, which is intrinsically dependent on disaster return times^2^. Third, whilst cereals have been the predominant focus of global analyses of climate impacts^3,7-9^, differences in production geographies^10^ as well as renewed focus on nutrition^^11^,12^ suggest a need to assess climate disaster risk profiles across different commodity groups.

Here we attempt to fill these three knowledge gaps by estimating the cumulative impact of nationally reported extreme heat and drought disasters occurring in 131 countries on the productivity of six major commodity groups (cereals, oilcrops, pulses, roots and tubers, vegetables and fruits), over the period of 1961-2014. Following previous work^3^, we utilize disaster occurrence data from the EM-DAT CRED International Disasters Database^13^, and crop production and value time series data from the United Nations Food and Agricultural Organisation^14^. We estimate national production deviations during heat and drought disaster years for each country and commodity compared to a counterfactual without disasters. We then use historical simulations to identify the null distribution of production deviations in each country in non-disaster years. This methodology provides new insights into the countries that show out of the ordinary crop production deviations in years in which extreme weather disasters were reported. In addition to calculating the impacts associated with heat and drought disasters, we also identify the global cost of these losses in monetary terms and the profile of monetary losses across all nations for which notable production deviations occurred.

Globally, we estimate that 1.4% of cereal production, 0.5% of oilcrops, 0.6% of pulses, 0.2% of fruits, and 0.09% of vegetable were lost due to heat and drought disasters over 1961-2014. Our improved estimate of global cereal production loss is almost half the previous estimate of the impact of heat and drought events^3^ which pooled counterfactuals across countries globally. This is the first time to our knowledge that the global crop production losses to heat and drought events for non-cereal commodity groups has been calculated.

Our results show substantial variation in national responses to heat and drought (Figure 1). The largest drag from heat and drought for cereals were observed in Botswana (5.7%), followed closely by the USA (4.4%), Australia (4.4%), and Angola (4.1%). There are a few things to note about these losses. First, these national losses deviate markedly from the global loss estimate. Second, there are some countries where the production losses under heat and drought events are close to the null distribution for these types of disasters (e.g. Botswana), and others where associated losses fall far outside the natural variation in production (e.g. Angola and USA; Figure 1A). This illustrates that the perception of heat and drought disaster risk is likely to greatly depend on the other factors that drive inter-annual production variation within each country. Third, our analyses show equal levels of long-term risk in percentage terms in both developed and less developed countries. Thus on a percentage basis, disaster risks might not be greater in technologically advanced farming systems as had previously been suggested^3,15^.

**Fig 1.**
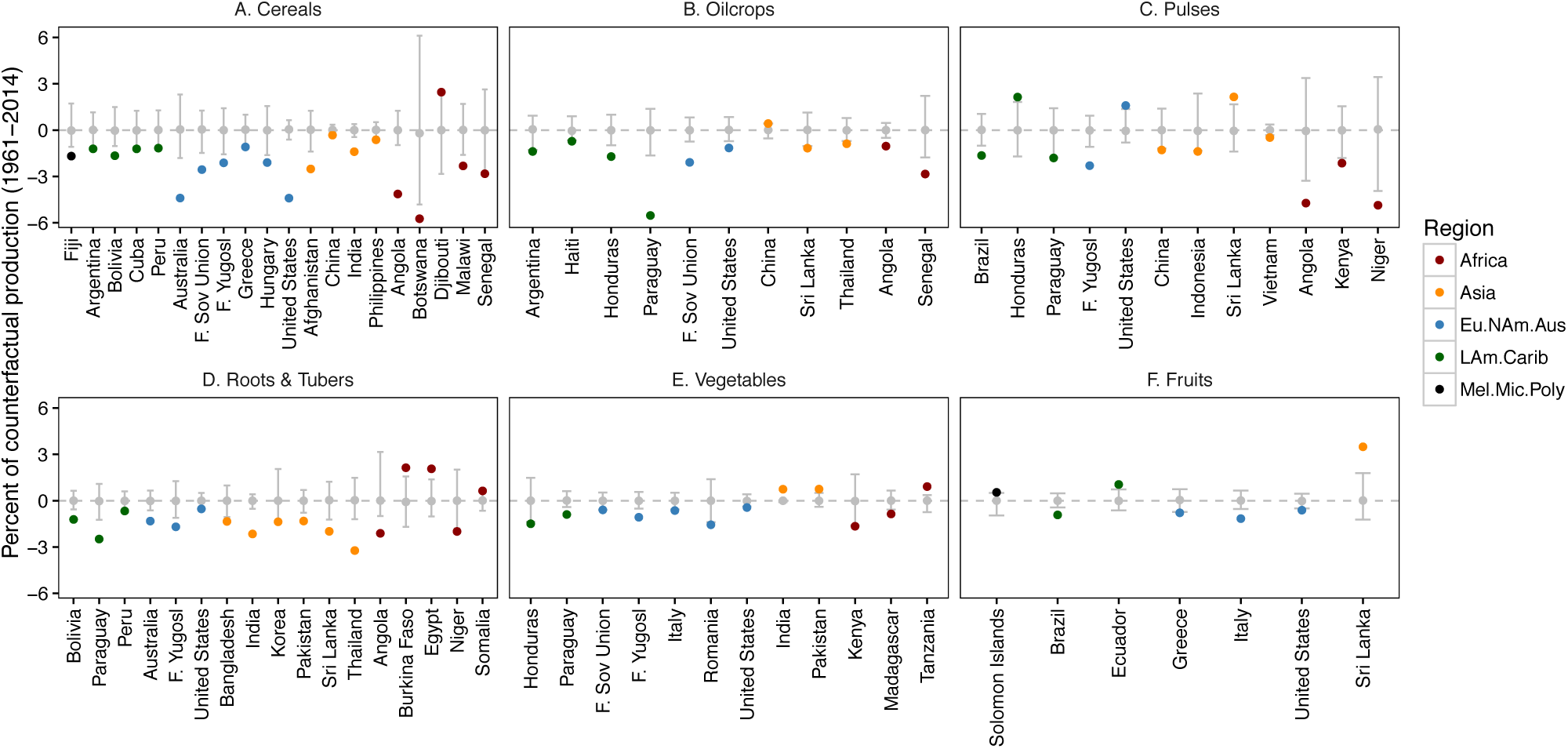
Effect of heat and drought disasters on global crop production. All the cases where a significant production loss or gain was estimated are shown. The y-axis indicates the percent of production within a country that was lost or gained during heat and drought events over 1961-2014. Gray points and whiskers show the median and range of the null distribution for losses or gains in years when heat and drought events did not occur.

Our analysis also shows substantial variation in the influence of heat and drought across different commodity types. The largest non-cereal losses occurred in Paraguay for oilcrop production (5.5%), Angola and Nigeria for pulse production (4.7% and 4.8% respectively), and Thailand for roots and tubers (3.2%). This commodity comparison provides two additional insights. First, there are differences between commodity responses within countries during heat and drought events. For example, over the study period the USA saw significant losses in cereal (see above) and oilcrops (1.1%), but significant gains in pulse production (1.6%). The presence of positive deviations, or lack of significant impacts in certain commodities where others fail, is suggestive that commodity diversity might offer some alleviation of risk to extreme events due to portfolio effects – a benefit of biodiversity well recognized in the ecological literature (see Ref 16). Secondly, some commodities seem more susceptible than others – the least severe losses occur in vegetables, fruits, and roots and tubers, and the most severe in cereals, oilcrops and pulses. These differences suggest that an assessment of sustainable diets^11,12^ might also benefit from identifying the ‘climate riskiness of the plate’ in addition to other environmental and social considerations.

In monetary terms we estimate the net effect of heat and drought events across all commodities and countries over the study period to be ~$237 billion USD (2004-2006 baseline). These losses were not evenly distributed across countries – the largest financial losses were incurred by USA ($116 billion), Soviet Union ($37 billion), India ($28 billion), China ($11 billion) and Australia ($8.5 billion) (Figure 2) (Although our estimates for China are more conservative than for other countries, see Supplementary Information). Losses were also not evenly distributed across commodity types, with the vast majority being due to cereals ($190 billion), and the remaining allocated to pulses ($3.4 billion), oilcrops ($19 billion), roots and tubers ($9.3 billion), fruits ($12 billion) and vegetables ($2.1 billion). These monetary impacts show substantial bias for losses toward countries holding the world’s major breadbaskets, and towards crops that make up most of human calorific intake. These figures highlight the potential economic opportunity from reducing vulnerability and exposure to extreme heat and drought events in arable agriculture.

**Fig 2.**
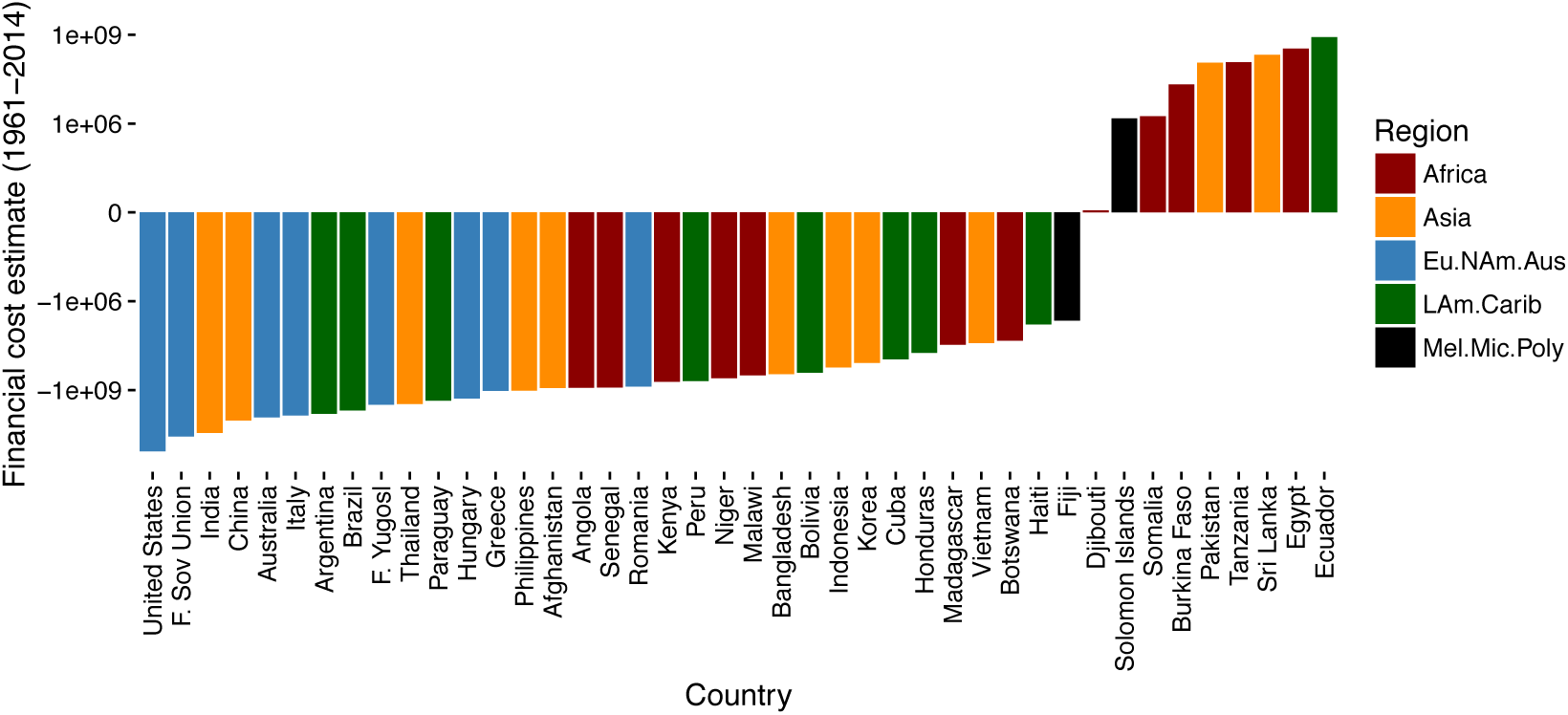
Estimated economic cost of heat and drought disasters on global crop production during 1961-2014. Losses and gains in production from figure 1 were converted into dollar values and summed for each country. A y-axis value of –1e+09 is equal to a loss of 1 billion USD (2004-2006 baseline). Note the logarithmic scale on the y-axis.

We have four key messages from this analysis. First, we find large variation in the effects of heat and drought disasters on crop production at the national level, which has to date gone unrecognized in global analyses. Second, we find evidence for significant drag on crop production in countries in Africa and Asia that on a percent basis equal those in the USA or Europe. This contradicts previous analysis that estimated regional averages and suggests that both developed countries and less developed countries can be equally susceptible on the national level to droughts and heat waves. Third, we observed differences between commodities in the historical impacts of heat and drought events. These differences between nations and commodities suggest that our risk profiles to extreme events will depend on what we choose to consume and in which country we choose to grow it. How future consumption trends influence the climate risk comprises an important avenue of future research. Finally, we found that the financial losses from extreme heat and drought events are not trivial and are not evenly allocated across countries. We show significant economic opportunity from avoiding similar losses to heat and drought events in future – particularly for large agricultural producers such as the USA.

In sum, our analysis provides the first global picture of cumulative losses associated with drought and heat events across different commodity types at the country level, and the first monetary evaluation of these losses. We hope that this work will help better integrate scientific assessments of risk into international disaster risk response profiling initiatives, will aid proactive action to prevent losses in the future, and garner support for designing more resilient global cropping systems.

## Acknowledgements

This research was supported by an NSERC Discovery Grant, and a Genome Canada/Genome BC Grant to N. Ramankutty.

## Materials and Methods

A full set of reproducible R^17^ script and data are supplied as Supplementary Information to enable others to undertake the entirety of the analysis presented in this paper. Here we provide a brief description of the analysis.

## Data sets

Three open source data sets were used in the analysis presented here. We obtained records of extreme weather disasters from the EM-DAT CRED International Disaster Database (http://emdat.be/), and crop production and gross production value data from the United Nations Food and Agricultural Organization’s FAOSTAT database (http://www.fao.org/faostat/en). We processed the data to maintain continuity in geographic boundaries over time (e.g., aggregating data from 1992 onward to the Former Soviet Union). We matched the production data to the countries and year of recording present in the disaster database.

## Disaster impacts

To identify the impact of heat and drought events on production for each of the six commodities for each country, we constructed a counter factual production in disaster years and compared it to the observed production in those years. To do this we created two complementary 3D arrays, *x*_*i* = 1:131, *j* = 1:53,*k*·*l* = 1:18_ and *y*_*i* = 1:131,*j* = 1:53,*k*·*l* = 1:18_ containing the national level production data for disaster and non-disaster years respectively, where *i* are countries, *j* are years (1961-2014), *k* are the commodities (cereals, oilcrops, pulses, roots and tubers, vegetables and fruits), and *l* are disasters (heat, drought, heat & drought). Counter factual production in disaster years was estimated by linearly interpolating between *y*_*j*_′*s* to create a new array *ȳ*_*i,j,k*·*l*_.The loss or gain during heat and drought events for each country and commodity, *L*_*i,k*_, was then estimated by summing the differences between the observed production and the counterfactual for all disaster types, *L*_*i,k*_ = **Σ**_*j,l*_ (*ȳ*_*i,j,k*·*l*_ – *x*_*i,j,k*·*l*_). The cumulative impact during heat and drought events for each country for each commodity, *I*_*i,k*_ was then estimated as *I*_*i,k*_ = *L*_*i,k*_ / (*P*_*i,k*_ + *L*_*i,k*_), where *P*_*i,k*_ is the sum of observed production for a given country and commodity over the study period, 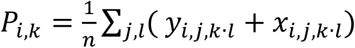, where *n* is the length of *l*. Thus, *I*_*i,k*_, identifies the percent loss or gain in crop production for a given country and commodity, over the study period, against a counter factual in which the disaster did not occur.

## Null distributions

To identify if the production deviations in *I*_*i,k*_ were no different from what would be expected in years in which heat or drought disasters did not occur, we calculated the null distributions for each element of *I*_*i,k*_, by running a 1000 simulated histories. We first set production values in real disasters years to null values. Then, for each of the 1000 simulations, we randomly generated three fake disaster occurrences to occur in each country (the median number of heat and drought disasters occurring over the 1961-2014 across countries was 4, and the range was 1-25). We used these fake disasters to create two more complementary 3D arrays, *xF*_*i* = 1:131,*j* = 1:53,*k* = 1:6_ and *yF*_*i* = 1:131,*j* = 1:53,*k* = 1:6_, containing the national level production data for fake disaster and non-disaster years respectively. Counter factual production for fake disaster years was estimated by linearly interpolating between *yF*_*j*_′*s* to create a new array 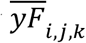. The loss or gain during fake disasters for each country and commodity, *LF*_*i,k*_, was then estimated by summing the differences between the observed production and the counterfactual, 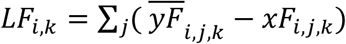. The cumulative impact of fake disasters for each country for each commodity, *IF*_*i,k*_ was then estimated as *IF*_*i,k*_ = *LF*_*i,k*_ / (*P*_*i,k*_ + *LF*_*i,k*_). *I*_*i,k*_ elements falling outside the bounds of the distribution of *IF*_*i,k*_ highlight the countries and commodities that show cumulative deviations in production during heat and drought years that are more extreme than deviations in years in which heat and drought events did not occur.

## Monetary impacts

To estimate the total value of crop production for each commodity and country in our analysis, *V*_*i,k*_, we retrieved the annual Gross Production Value (constant 2004-2006 terms) for each of commodity and country, and summed these for the years 1961-2014. To estimate the cost of heat and drought events for each country and commodity, *C*_*i,k*_ we then multiplied the values of production, by the percent loss or gain in crop production for a given country and commodity against the counter factual *C*_*i,k*_ = *I*_*i,k*_ · *V*_*i,k*_.Thus, *C*_*i,k*_ indicates the dollar value of production that might have been obtained if heat and drought events did not occur for a given country and commodity, under the assumption of linear pricing with respect to supply. We summed over all commodities, *k*, to estimate the net impact of heat and drought events on a country basis.

